# 3-D Reconstruction of Fingertip Deformation during Contact Initiation

**DOI:** 10.1101/2025.02.10.637366

**Authors:** Donatien Doumont, Anika R Kao, Julien Lambert, François Wielant, Gregory J Gerling, Benoit P Delhaye, Philippe Lefèvre

## Abstract

Dexterous manipulations rely on tactile feedback from the fingertips, which provides crucial information about contact events, object geometry, interaction forces, friction, and more. Accurately measuring skin deformations during tactile interactions can shed light on the mechanics behind such feedback. To address this, we developed a novel setup using 3-D digital image correlation (DIC) to both reconstruct the bulk deformation and local surface skin deformation of the fingertip under natural loading conditions. Here, we studied the local spatiotemporal evolution of the skin surface during contact initiation. We showed that, as soon as contact occurs, the skin surface deforms very rapidly and exhibits high compliance at low forces (<0.05 N). As loading and thus the contact area increases, a localized deformation front forms just ahead of the moving contact boundary. Consequently, substantial deformation extending beyond the contact interface was observed, with maximal amplitudes ranging from 5% to 10% at 5 N, close to the border of the contact. Furthermore, we found that friction influences the partial slip caused by these deformations during contact initiation, as previously suggested. Our setup provides a powerful tool to get new insights into the mechanics of touch and opens avenues for a deeper understanding of tactile afferent encoding.

## Introduction

Our remarkable ability to dexterously manipulate objects, such as writing with a pen, buttoning a shirt or fetching a key in a pocket, essentially originates from the use of our fingertips (Johansson and Flanagan 2009). Human fingertips serve as fundamental sensory units for effective object manipulation (Johansson and Westling 1984), utilizing feedback from thousands of mechanoreceptors embedded within the skin and distributed across its surface (Corniani and Saal 2020). When mechanically stimulated, these receptors sense skin deformations and in turn send precise spiking responses to the central nervous system (CNS) (Johansson and Vallbo 1979; B. B. Edin 1992; Benoni B. Edin 2004; Jarocka et al. 2021). As such, fingertips provide rich information about high-level features such as object geometry, orientation, and slipping state, enabling feedback that helps fine-tune gripping behavior (Johansson and Westling 1984; LaMotte and Srinivasan 1993; Jenmalm et al. 2003; Goodwin and Wheat 2004; Witney et al. 2004; Saal et al. 2009; Schiltz et al. 2022).

Neurophysiological recordings in both humans and non-human primates have revealed that most tactile afferents innervating the fingertip, including those located far from the point of contact, are highly active during normal and/or tangential loading (Khalsa et al. 1998; Birznieks et al. 2001; Birznieks et al. 2010; Jenmalm et al. 2003; Goodwin and Wheat 2004). This finding suggests that significant skin deformation extends well beyond the contact area during mechanical stimulation. These deformations are all precisely encoded and contain rich information about the dynamics of the stimulation. Quantifying them is essential to uncover the richness and diversity of tactile afferent feedback during contact interactions with objects (Deflorio et al. 2022). Yet, our understanding of the mechanical response of fingertip skin to even simple interactions — at the stimulus site and at a distance — remains very limited (B. Delhaye et al. 2016). As a result, while essential features of tactile stimulation can be readily decoded from the response of population of afferents (B. P. Delhaye et al. 2019; Khamis et al. 2015), it remains virtually impossible to predict individual responses in natural conditions of manipulation, that is, with shear forces and normal component above 1 N (for other cases see Saal et al. (2017)).

To address this, two opposing approaches have been suggested by Vincent Hayward (Hayward et al. 2014; Jörntell et al. 2014): The first consists in delivering a precisely controlled, localized stimulus with well-characterized skin deformation dynamics — for example, using a device that he developed called the Latero Tactile Stimulator (Hayward and Cruz-Hernández 2000). The second is to directly measure skin deformation as a consequence of a given stimulus, for which he pioneered the imaging approach to track fingerprint deformation (Levesque and Hayward 2003), which largely inspired the present study.

*In vivo* studies have measured fingertip skin surface deformation in both passive and active experiments by tracking fingerprint motion using an optical arrangement called frustrated total internal reflection FTIR (B. Delhaye et al. 2016; Willemet et al. 2021; Schiltz et al. 2022; Dunilac et al. 2023). However, these methods are inherently constrained to capturing deformation within the contact area (planar). Other techniques, such as optical coherence tomography (OCT), have shown promise in imaging sub-surface deformation near the depth of mechanoreceptors, achieving high spatial resolution of the order of microns. Still, they are currently restricted to measuring only a few millimeters of cross-section (Corniani et al. 2024). As such, computational models are valuable tools for predicting and analyzing skin deformation when direct measurements are challenging. However, given the complex nature of fingertip skin – multilayered, hyperelastic, viscoelastic, and anisotropic (Pawluk and Howe 1999; Groves et al. 2013; Duprez et al. 2024), there is a need for precise and full-scale measurements of fingertip skin responses to inform and validate these models. Recently, Kao et al. (2022) applied ink to the skin surface and combined stereovision with digital image correlation (DIC) to measure fingertip skin deformation in response to light and localized stimulation near the perceptual threshold, using von Frey filaments. This method enabled the detailed reconstruction of the fingertip surface while minimally affecting its biomechanics.

Building on fingerprint imaging and the DIC approach, the present study extends previous empirical measurements by using multiple cameras to reconstruct the entire fingertip surface during typical human contact initiation against a flat, rigid surface. This approach allows us to measure the global response of the fingertip as well as the localized deformation response of the skin surface (accurate to within a fingerprint ridge). First, we provide a detailed description of our method, validated through multiple approaches, including comparisons with independent measurements. Next, we present evidence of significant deformation occurring both inside and outside the contact area. Notably, we observe strain waves propagating from the initial point of contact to the periphery in the meridional direction, closely following the contact border as the loading increases. Additionally, we demonstrate that friction, a critical parameter for object manipulation, influences partial slip occurring during the initial loading phase of the fingertip. This study aims to offer a more comprehensive and quantitative understanding of fingertip skin dynamics during contact initiation, providing both global and localized biomechanical responses.

## Materials and methods

### Participants

Nine volunteers (two female, 29 *±* 5 years of age, mean *±* SD) participated in the experiments. They all provided written informed consent and the experiment was approved by the local ethics committee.

### Robotic platform

A custom robotic platform was used to apply controlled stimuli to the fingertip (Fig. 1A). A detailed description of this system has been previously published (B. Delhaye et al. 2014; B. Delhaye et al. 2016; Dunilac et al. 2023). In short, a transparent plate was mounted horizontally on the end effector of a 4-DOF (degrees of freedom) industrial robot (Denso HS-4535G, Denso Robotics, Japan) and coupled with two force/torque transducers positioned on each side of the plate (Mini 40, ATI Industrial Automation, USA). The robot controlled the plate’s lateral motion in the horizontal plane. The participant’s right index finger was attached to a support designed to guide the nail and control the finger’s motion in the vertical plane. This support was driven by a servomotor (Dynamixel XM430, ROBOTIS Co., Ltd., Korea) with its axis of rotation closely aligned to the participant’s metacarpophalangeal joint, thereby mimicking the natural motion of the finger. The control law implemented used real-time normal force feedback from the plate to adjust the servomotor’s position, achieving a target normal force of 5N at a defined constant loading speed (see Experimental procedures). The angle of attack between the longitudinal axis of the finger and the plate was maintained within 20-25deg.

**Figure 1.**
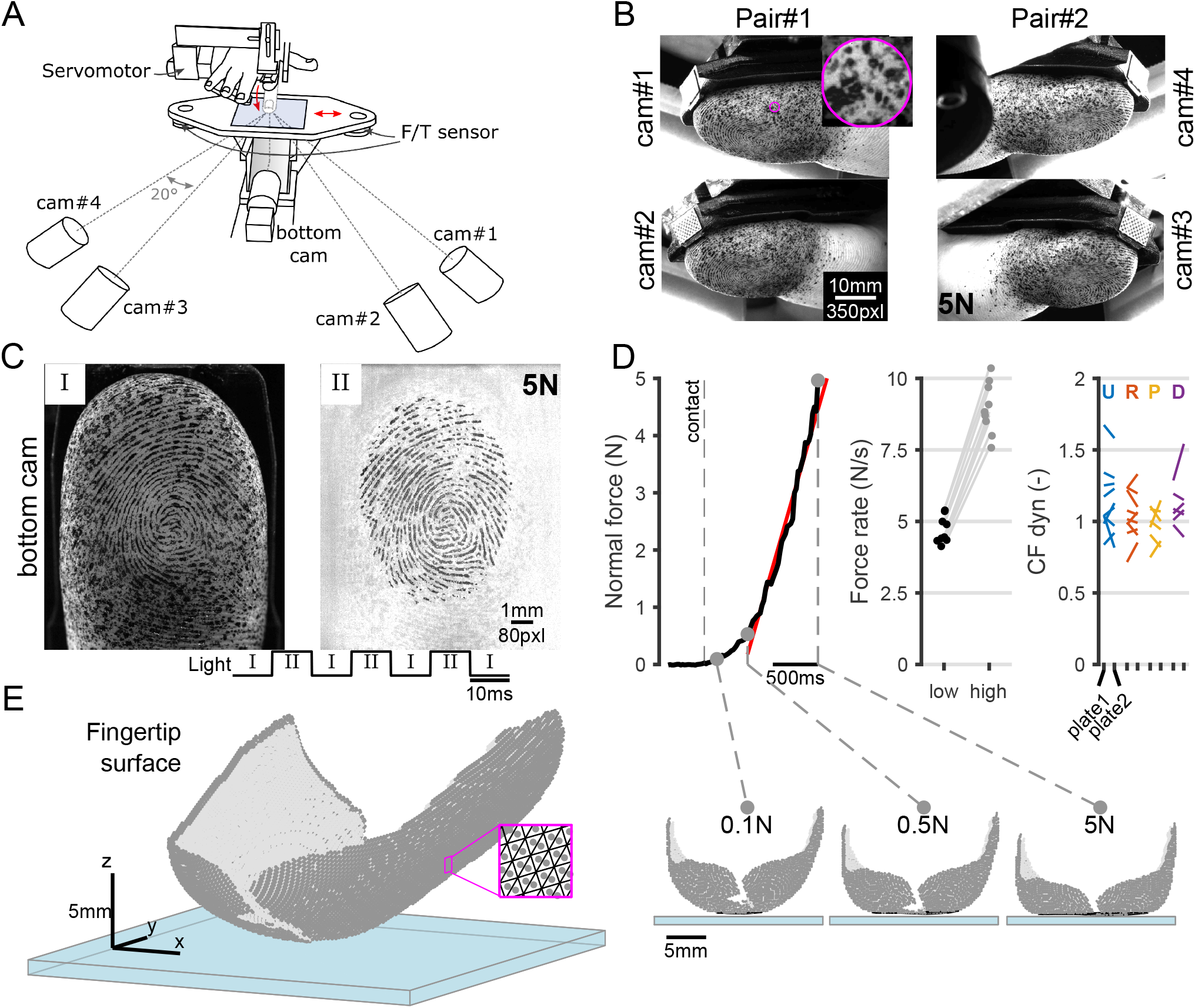
Experimental setup, protocol, and reconstruction examples. (**A**) The right index finger of participants is guided via a servomotor to stroke a transparent plate attached to a robotic platform, with an imaging system comprised of five cameras capturing a portion of the fingerpad. (**B**) Typical images (S02) from each stereopair of cameras used for 3D reconstruction during normal loading are shown at 5 N, along with a zoomed-in view of the speckle pattern from cam#1. (**C**) Pair of images captured by the bottom-facing camera at a 10 ms interval with alternating light sources. Images from (B) are taken in the “I-phase”. (**D**) Left: One typical normal force trace during fingertip loading. Center: Force rates for each loading *Speed* condition, calculated from a linear fit (red segment in (D, left)) of the force signal between 0.5 and 4.5N (n = 9). Right: Dynamic coefficient of friction evaluated during tangential loading for two plate conditions (hydrophilic and hydrophobic) and four tangential directions (U = ulnar, R = radial, P = proximal and D = distal). Median values for each subject are shown. (**E**) 3D reconstruction of the fingertip surface from an oblique projection view (left), and front projection view (right) at different loading stages

### Imaging apparatus

The robotic platform was combined with an imaging apparatus composed of an array of four cameras forming two stereo pairs (Fig. 1A,B, Suppl. Movie M1), synchronously triggered at 50 frames per second (fps) (2MP, ALVIUM 1800 U, and 50mm-focal lens Fujinon CF50HA) situated around the fingertip, and a bottom-facing camera triggered at 100 fps (Fig. 1A,C, as in Dunilac et al. (2023), 5MP, JAI-GO-5000M-PMCL and 50mm-focal lens Ricoh FL-BC5024-9M). Polarizing filters were added to the camera array lenses to reduce specular reflections, which significantly improved the imaging quality. After calibration, the camera array and the bottom camera achieved image resolutions of approximately 35-40 pxl/mm (depending on the scene depth) and 80 pxl/mm, respectively.

The camera array placement was carefully designed based on the fingertip’s geometry and the optical characteristics of the camera lens. First, to maximize the extent of the reconstructed surface of the fingertip and capture deformation both inside and outside the fingertip-plate interface. Second, to limit the projective distortion while keeping appropriate out-of-plane precision. Hence, each stereo pair was placed around 45° from the longitudinal axis of the finger on each side of the index and inclined by 30° below the horizontal. Stereo angles for DIC are often prescribed between 15 to 30°, as this range provides sufficient overlap and limits distortion while ensuring accurate out-of-plane displacement measurements. In this setup, a stereo angle of approximately 20° was used. The imaging apparatus included two different light sources that illuminated the fingertip alternately at 100 Hz (see bottom of Fig. 1C). First, a diffused light source (LED strips with diffusing material) was positioned around the fingertip to create uniform illumination across its surface. Second, a light source coaxial to the bottom-facing camera directed light through a semi-transparent mirror onto the fingertip. The latter leveraged the principle of frustrated total internal reflection (FTIR) to create high contrast between fingerprint ridges and valleys (B. Delhaye et al. 2014). Synchronization of the imaging setup was achieved through a TTL (transistor to transistor logic) signal sent by the robotic platform to the external trigger inputs of the cameras and light sources.

### Imaging approach

We adopted an imaging approach called stereo digital image correlation (or 3D-DIC) to measure the dynamic response of the fingertip under load. 3D-DIC is a non-contact, full-field measurement technique that captures the three-dimensional surface geometry and displacements of a deformed surface. In short, this technique establishes a correspondence between a reference frame and subsequent frames by dividing the reference frame into small sections, known as subsets, and locating these subsets in subsequent frames (2D-DIC). To transition from 2D to 3D, subset locations are matched across a stereo pair of images and mapped into 3D space through calibration. A comprehensive description of the methodological aspects of 3D-DIC can be found in the literature (Sutton et al. 2009). Specifically, we used the open-source 3D-DIC toolbox, MultiDIC (Solav et al. 2018), developed in MATLAB and built on top of Ncorr, a 2D-DIC algorithm (Blaber et al. 2015). This toolbox was adapted to account for large displacements, as explained in the paragraph below. For each trial and participant, the reconstructed surface yielded approximately 15,000 triangular elements tracked over 150 time steps (Fig. 1E). At each element, the Lagrangian finite strain tensor E was computed, from which were derived the principal Lagrangian strain components (E1 & E2) and maximum shear strain, as described in MultiDIC. These finite strain measures are typically employed for analyzing biological soft tissues undergoing large deformations.

To improve tracking accuracy, we adapted the 2D-DIC step by segmenting it into three sequential steps within each stereo-pair, a method called partitioned correlation (Jones and Iadicola 2018). This modification ensured robust image tracking of material points by assigning a distinct reference frame to each 2D-DIC process, thereby reducing error accumulation (similarly to the strategy in Pan et al. (2012)). More specifically, the first step consisted of mapping the material points of the initial stereo-pair frames (Fig. 2A-B, M for “matching”, Suppl. Movie M2-3). Using the first frame of each camera as the reference, material points were then tracked over time in subsequent frames (Fig. 2A-B, T1 and T2 for “tracking” on cam#1 and cam#2). This approach accounted for not only fingertip deformation but also the distortion introduced by the curved geometry of the fingertip and varying angular projection views from the cameras.

**Figure 2.**
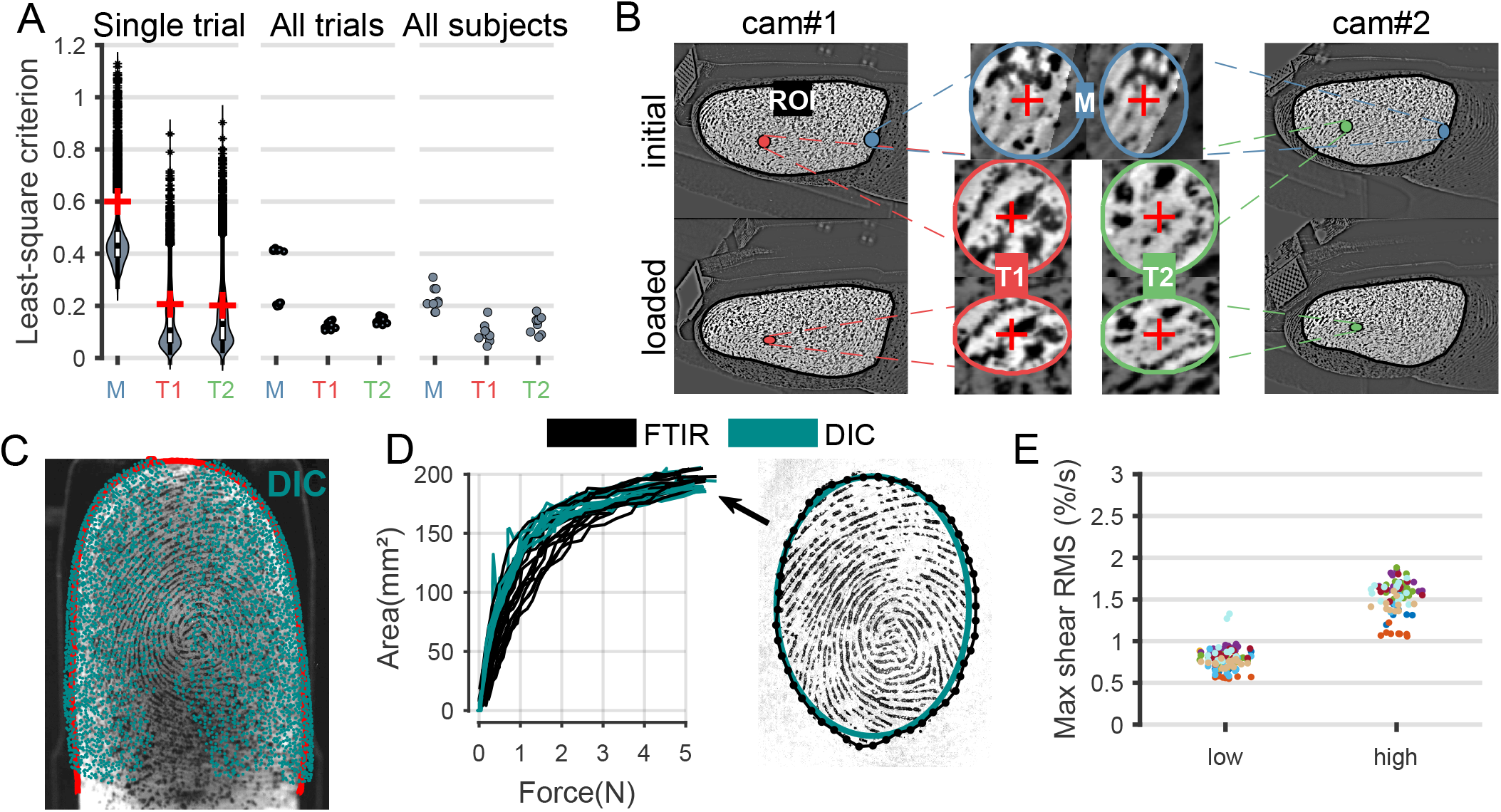
Validation of the reconstruction method from low-level to high-level metrics. (**A**) Least-squared criterion distribution for one stereo-pair in a single trial, median values for all trials for one subject, and median across all trials for all subjects. (**B**) 2D-DIC tracking evaluation within a stereo-pair illustrated using three example subsets. Subsets are displayed in both the reference (initial) and deformed (loaded) configuration for each correspondence procedure: matching (blue subset), tracking 1 (red subset), and tracking 2 (green subset). Least-squared criterion levels are represented by a red cross in (A, left). Subset size is accurately depicted. (**C**) Comparison of fingertip geometry between the bottom camera and the 3D-DIC reconstruction made before contact initiation. The bottom camera image contour is shown in red, while the DIC reconstruction projected from a below view is overlaid in blue. (**D**) Evaluation of contact area contours using two methods: DIC (fitted ellipse from point cloud) and FTIR images. The evolution of contact area with increasing loading force is shown for a single subject with 14 repetitions. (**E**) Noise levels evaluated as RMS of maximum shear strain rate before contact for both speed conditions across all trials, with each color representing an individual subject.

### Experimental procedures

Before each experiment, the index fingertip was patterned using ink to achieve a high signal-to-noise ratio (SNR) for digital image correlation, as the natural fingerprint pattern lacked sufficient features for tracking. Following the procedure described by Kao et al. (2022) (without using a layer of paint), a dense, random, texturized speckle pattern was created on the fingertip surface. This pattern adhered to the skin and deformed with it, facilitating accurate tracking. The speckle pattern was created by spraying ink droplets onto the fingertip using an air compressor (Master AirBrush, TCP Investments, LLC, Florida, USA). The speckles had an average diameter of 250-300 µm (*∼*10 pixels) and covered around 40% of the measured skin surface. Due to friction during contact, some ink in intimate contact with the glass plate wore off, requiring the speckle pattern to be reapplied twice: once before the first trial and again midway through the experiment. After the experiments, the ink was removed with soap and water.

The fingerpad was loaded under conditions mimicking the initial instant of grasping (Forssberg et al. (1991), see Fig. 1D). In each trial, the transparent plate was centered under the fingertip. While the plate was kept still, the fingertip was loaded onto the plate at either 3.5 deg/s or 7 deg/s (*∼*5 mm/s or 10 mm/s) until stabilizing at a normal force of 5 N (pseudo-randomized trials). These loading speeds resulted in force rates of 4.45 *±*0.92N/s (95% CI) and 8.83 *±* 1.93N/s (see Fig. 1D), correspond to low- and high-speed conditions, with a force rate ratio of 1.93 *±* 0.125. After achieving the 5 N normal force, the plate was moved laterally (horizontally) at a constant speed of 5 mm/s in one of four directions: Ulnar, Radial, Distal, or Proximal. This motion caused the fingertip to reach full slip, with a total lateral displacement of 16 mm. While data were collected during this period, only the frictional properties of the contact were analyzed in this study (see *Data analysis*).

Two types of glass plates were used: untreated hydrophilic glass and chemically treated hydrophobic glass. The hydrophobic treatment was intended to alter the friction coefficient while maintaining a flat, transparent surface. Although preliminary tests indicated that the treatment slightly lowered friction, the ink on the fingerpad interfered with this effect, leading to inconsistent behavior across participants and canceling any significant trends (Fig. 1D right). Each loading condition was repeated eight times, for a total number of 32 trials of normal loading per participant, evenly distributed across sliding conditions. Before each trial, the plate was cleaned thoroughly with alcohol to remove any sweat residues.

### Membrane experimental model

In addition to reconstructing the fingertip of human participants, we applied the same reconstruction procedure to measure the loading response of an air-filled latex membrane. A balloon created from a latex finger glove was strapped to the nail support of the robotic platform and subjected to the same loading conditions as the fingertip. The diameter of the balloon (28 cm) was close to the fingertip size, although it was not further calibrated to replicate the elasticity of fingertip skin. While the coefficient of friction between the latex balloon and the glass plate was not measured, it is reasonable to assume a high friction level, consistent with findings on latex-glass interactions (Carré et al. 2017). The empirical data obtained from the balloon were compared to predictions from an analytical model of an inflated spherical hyperelastic membrane pressed against a flat rigid surface (Kumar and Dasgupta 2013). In contrast with Hertzian contact mechanics, this model does not rely on the assumption of small strains, the violation of which can lead to significant prediction errors (Dintwa et al. 2008). Additionally, it accounts for the presence of contact friction. While the membrane model is highly simplistic compared to the complex geometry and structure of the human fingertip, it offers a useful framework for evaluating surface strains under large deformations in the presence of frictional contact.

### Data analysis

All data analyses were implemented in MATLAB 2021a. For the first three subjects, approximately half of the trials were excluded due to visual obstructions caused by the robotic platform’s initial positioning. Design modifications subsequently resolved this issue, enabling full analysis of all trials for the remaining six subjects. Force data were low-pass filtered with a fourth-order digital Butterworth filter with a cut-off frequency of 40 Hz and zero phase lag. Surface strains were time-processed with the same filter at a 10 Hz cut-off frequency (1/5th of the image acquisition rate of 50 frames/s) and spatially filtered using a Gaussian filter (std = 0.5 mm) over a neighborhood area. The Gaussian filter width was chosen to span no more than two fingerprint ridges.

For the DIC process, material points within the region of interest (ROI) of the reference frame were tracked every 10 pixels, with circular subset radii of 40 pixels used for the tracking steps (Tracking 1 and 2) and a 60-pixel radius for the matching step (Fig. 2B). The increased radius for the matching step was necessary due to the greater challenge of correspondence in this phase of the DIC process. These radii were chosen to avoid excessive smoothing while ensuring effective tracking of the entire ROI. The spacing between tracked points was approximately 0.3 mm, which is around half the average ridge-to-ridge spacing of a fingertip. The criterion of correlation for the DIC procedure was expressed as a normalized least-squared criterion C_*LS*_, calculated for each subset S between the reference grayscale image f and the current grayscale image g:

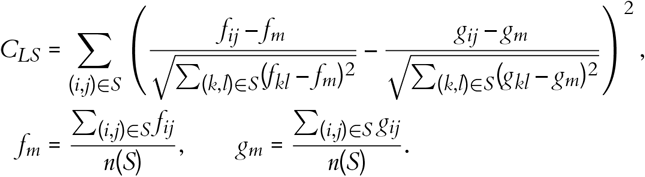

Subsets with low correlation values (C_*LS*_ > 1) were removed from the 3D measurements. Low correlation could result from visual occlusions, specular reflections, insufficient or excessive ink application, or out-of-focus images. Additionally, error-prone points at the border of the reconstructed stereo pairs were removed, which sometimes created the appearance of fractures along the borders of the reconstructed surface (Fig. 1E).

#### Friction computation

The dynamic coefficient of friction was calculated as the ratio of tangential force to normal force, averaged during a full-slip phase of each trial. This phase was conveniently defined as beginning after 10 mm of lateral displacement of the platform, ensuring that full slip had occurred, as confirmed by the characteristic plateau in tangential force following full slippage (B. Delhaye et al. 2014).

#### Gross contact area

The gross contact area was obtained using two independent methods. The first method used FTIR images captured by the bottom-facing camera, following previously described methods (B. Delhaye et al. 2014). The second method relied on 3D-DIC data, where the gross contact area was determined through two proxies. During the initial loading phase, points were considered in contact if their vertical speed fell below a threshold. This threshold was adjusted such that the resulting area would best fit the area measured with FTIR measures. As the contact area expanded and fingertip deformation reached saturation, a horizontal plane was fitted to approximate the plate position, and points at the vicinity of this plane were classified as being in contact. For consistency, the gross contact area is referred to as the “contact area” throughout the paper.

#### Spatial distribution of deformation

The analysis focused on the distal region of the fingertip, where resolution was highest and distortion was lowest due to the camera arrangement and fingertip geometry. This region also approximates a half-spherical shape, facilitating comparisons with the membrane model and the latex balloon. To study the spatial distribution of deformation along the meridional direction, a normalized segmentation of the fingertip was defined for each participant at the 5 N loading instant. Inside the contact border, multiple concentric annuli were defined as fractions of the radii of a fitted ellipse representing the contact area. Outside the contact border, regions were segmented vertically, starting from the fitted plane of contact and evenly spaced by the same interval as the annuli inside the contact. Deformations were circumferentially averaged along each ellipse to examine their progression along the meridional direction.

### Statistical analysis

A linear mixed-effects model was used to study the effects of loading speed (*Speed*) and fingertip width (*Width*) on two dependent variables: contact area (*CA*) and deformation (*Def*). Mixed-effects models are well-suited for accounting for between subject variability by allowing random variations in the model’s intercept and slope. In this analysis, *Speed* was considered a fixed effect, while Subject was considered a random effect that could affect the model’s intercept. Similarly, the effects of repetition (*Rep*) and *Speed* on noise levels were analyzed using a similar framework. To investigate the effect of friction during loading, we took advantage of the variability in friction observed across trials for each subject, despite the lack of a significant influence from surface material (Fig. 1D right). For each trial, a mean slip value was calculated by averaging the displacements of points identified as being in contact. The effects of the coefficient of dynamic friction (*CFdyn*), *Speed*), and lateral direction (*Dir*) – from which *CFdyn* was derived – on the mean slip value were analyzed using a linear mixed-effects model. In this model, *CFdyn, Speed*, and *Dir* were considered fixed effects, while Subject was considered a random effect, allowing it to influence both the intercept and slope of the model. To test the significance of each fixed-effect term on the model results, an F-test was used.

## Results

### Experimental validation of the measurements

To ensure robust measurements, the methods were validated in four distinct ways: (1) assessing the DIC correlation levels alongside a visual inspection of the corresponding tracked features, (2) comparing our fingertip reconstructions with reference images taken from the bottom-facing camera, (3) estimating the noise level in the measured deformations, and (4) comparing the reconstructed response of a hyperelastic membrane using our method with that predicted by an analytical model (detailed in the next section).

First, we assessed the DIC criteria (see Material and Methods) used to correlate subsets of images in the three separate correlation processes detailed in the Imaging approach section: matching (“M”) and tracking (“T1” and “T2”) (Fig. 2A, left). Performance was analyzed at the single trial level, and the average least-squared criterion was computed for all trials and participants (Fig. 2A, center and right). To contextualize the least-squared criterion levels, Fig. 2B presents an illustrative example of a subset tracked during each correspondence procedure (matching and tracking) for three levels of least-squared criterion that are relatively poor compared to our average results (0.6, 0.2, and 0.2 respectively). These results demonstrate that the observed correlation levels were sufficient for effective tracking in our experiments, with similar levels consistently observed across trials and participants. Furthermore, the root mean square (RMS) error of the calibration 0.025 mm, approximately equivalent to the pixel level.

Next, we qualitatively compared the 3D-DIC reconstruction with 2D measurements obtained from the bottom-facing camera. By overlaying images from the bottom camera with the reconstruction projected onto the X-Y plane, we confirmed that the reconstructed fingertip geometry accurately matched the observed images (Fig. 2C). Additionally, we used FTIR images from the same bottom camera to segment the contact area, employing the method from B. Delhaye et al. (2014). The contact areas obtained from the reconstruction and the FTIR method showed excellent agreement, with a Spearman’s correlation coefficient of ρ = 0.945 (Fig. 2D). These measurements were also consistent with values reported in the literature (Serina et al. 1997; Dzidek et al. 2017).

Finally, we estimated the noise level in our measurements. Since no deformation should occur on the reconstructed fingertip surface before contact, noise was quantified by calculating the median shear strain rate during pre-contact periods across several repetitions. The noise was found to be approximately 0.8 %/s at low speeds and 1.6 %/s at high speeds (Fig. 2E), which is less than one-tenth of the peak signal amplitude observed during loading (see next section, Fig. 5B-C). Noise levels were strongly affected by *Speed* (estimate = 0.74; t_199_ = 0.83; p = 1e-118), but not by trial repetition (t_199_ = −1.83; p=0.41).

### Experimental validation with a model

To further validate our experimental approach, we compared the data with an analytical model describing the response of a hyperelastic balloon under normal load (Kumar and Dasgupta 2013). We assessed whether the observed deformation patterns– both in orientation and amplitude – were consistent with theoretical predictions.

#### Direction

The membrane model was formulated as an axisymmetric problem, with deformations expressed along the meridional direction (Fig. 3A, top). When the membrane was pressed against a flat surface, the model predicted that the principal deformations would align along the meridional and circumferential directions (Fig. 3A, bottom). Our experimental reconstructions revealed orientations of the principal strains (E1 & E2) that closely match the model, with a relative deviation of −13.1 *±* 38.35 deg (median *±* 95% CI).

**Figure 3.**
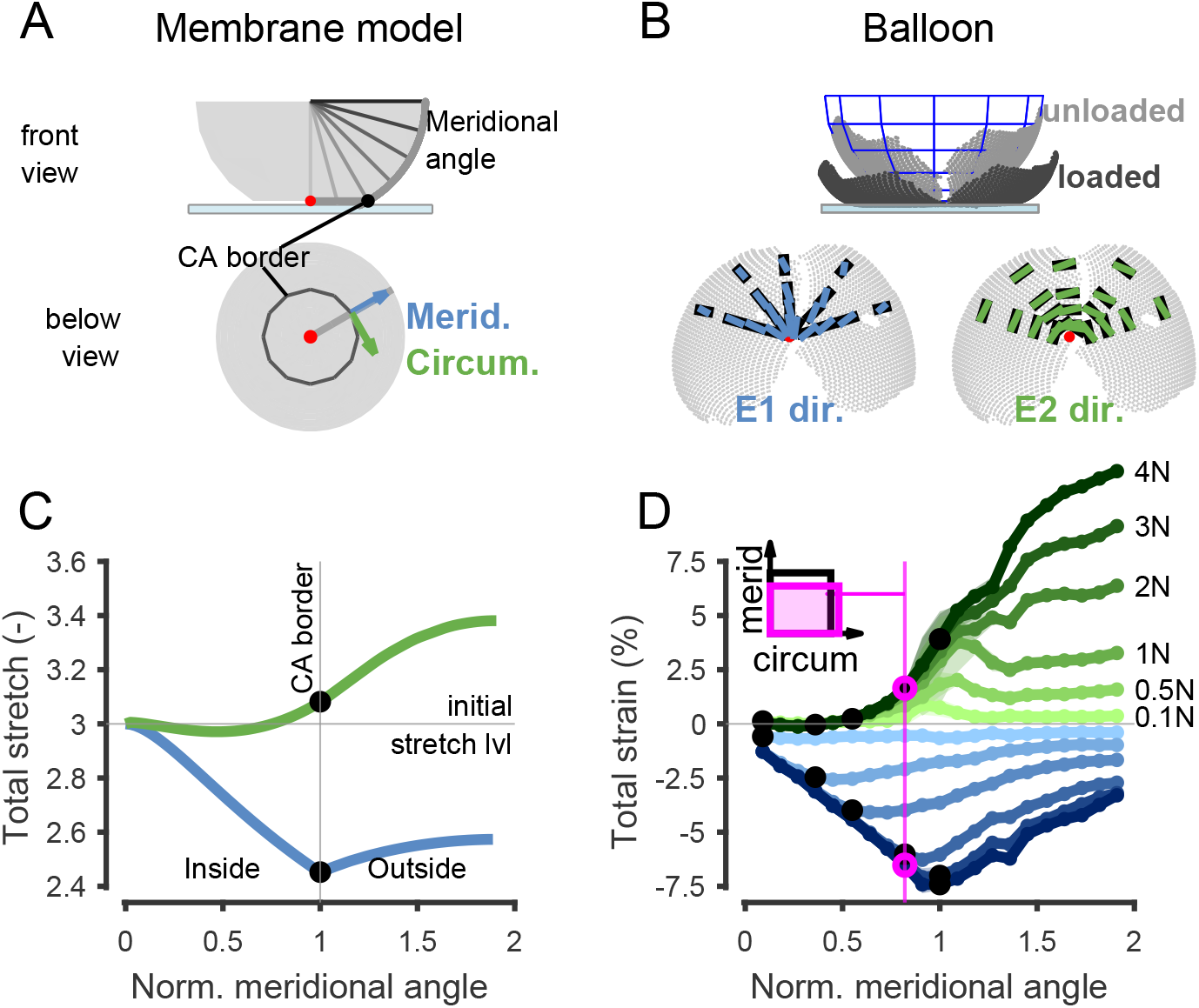
Analytical membrane model (data from Kumar and DasGupta 2013) vs experimental data of a balloon. (**A**) Membrane model principal directions (Below) in view from below and schematic of meridional angle in a front view (Top). (**B**) Balloon reconstruction. (Top) unloaded and loaded on the contact surface. (Below) principal directions (blue and green), and true meridional/circumferential (black). (**C**) Membrane model stretch evolution along a normalized meridional angle (below 1 = inside the contact /above 1 = outside the contact). (**D**) Balloon total strain evolution as in (C) at multiple loading steps. In the top left is an illustration of deformation values in meridional and circumferential directions (magnified by 3).

#### Amplitude

To enable comparisons between the theoretical model, the experimental balloon data, and the fingertip, we normalized the meridional angle so that the contact border has a value of 1. The model response (Fig. 3C) indicated that the first principal stretch is negative, while the second principal stretch is globally positive across the membrane periphery. This pattern reflects meridional contraction and circumferential elongation relative to the membrane’s initial stretched state (see reference for details). Interestingly, a peak contraction occurs at the contact border, sharply delineating the region inside the contact from the region outside. Meanwhile, the second principal stretch increases, reaching a maximum at the periphery of the membrane, outside the contact region. The empirical reconstruction of the balloon aligns well with the membrane model, exhibiting similar deformation trends under progressively increasing loads. Specifically, a contraction peak is observed at the contact border in the meridional direction, while positive strains in the circumferential direction increase to a peak value at the periphery of the balloon. As the load increases, high friction causes the contact area to stick to the plate, preventing further deformation within the contact region. Meanwhile, the area outside the contact remains free to move, leading to increasing strains in both principal directions outside the contact area as the load increases. It is important to note that the experimental parameters for the balloon are model were not identical, so absolute amplitudes of deformation should not be compared directly. However, the qualitative agreement between the observed and predicted deformation patterns reinforces the reliability of our experimental measurements.

### Fingertip response to normal loading

Similar to the balloon model, the fingertip exhibited significant shape changes and conformed to the planar surface under normal loading. The fingertip skin undergoes meridional contraction and circumferential elongation in the distal (tip) region (Fig. 4A, relative angular deviation 4.8 ± 35.3 deg; median *±* 95% CI). The principal strains (E1 & E2) increase with loading, reaching absolute amplitudes of approximately 5% to 8% on average at load of 5N (Fig. 4B). Interestingly, at forces as low as 0.05N, significant meridional contraction rates are observed, peaking at 20 % per sec. This forms a deformation front that shifts outward, closely following the contact border as the loading increases (Fig. 4C, blue). Similarly, circumferential elongation rates occur just outside the contact border (Fig. 4C, green). The deformation rates were relatively uniform along the circumferential direction, except for the later stages of loading, where deformation rates became more pronounced towards the distal fingertip (Fig. 4D-E, Suppl. Movie M4). Interestingly, the peak strain rate tends to be located outside the contact area, where the fingertip curvature is highest.

**Figure 4.**
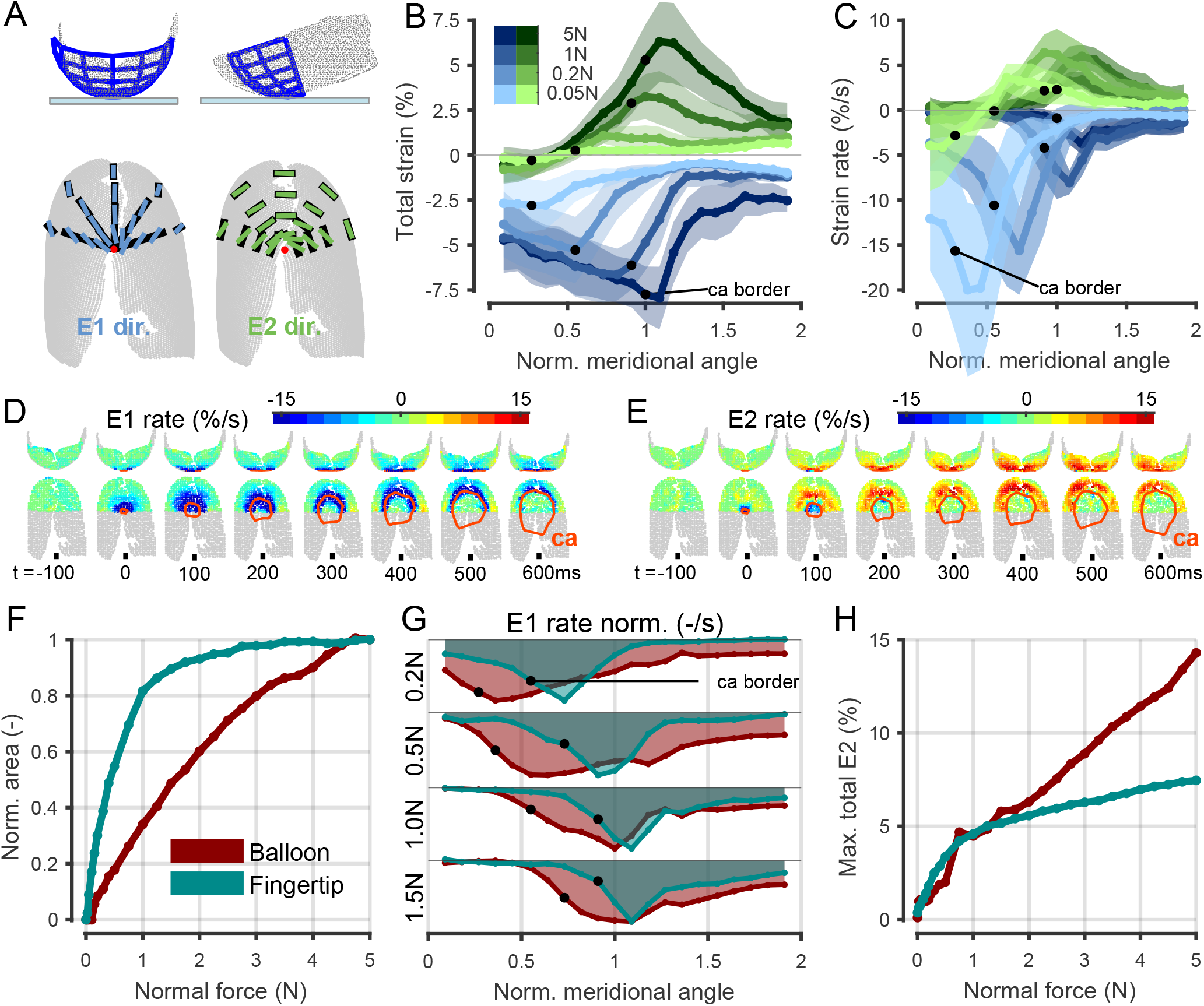
Fingertip mechanical response to loading. (**A**) Fingertip reconstruction. (Top) unloaded on the contact surface with a spherical fit in blue on the distal part of the fingertip. (Below) Principal directions (blue and green), and true meridional/circumferential (black). (**B**) Fingertip strain evolution during low-speed condition along a normalized meridional angle at multiple loading steps with contact border highlighted (black dot) and standard deviation across individuals (shaded regions). (**C**) Fingertip strain rate evolution (as in B). (**D, E**) Typical strain rate maps before and after loading initiation with contact border highlighted (ca). (**F**) Comparison between normalized contact area w.r.t. 5N versus normal force applied, for the balloon and the fingertip (median value between individuals). (**G**) Localization and extent of meridional strain rate along the surface at different normal force, for the balloon and the fingertip. Each trace was normalized to its peak value. (**H**) Max circumferential strain (95^*th*^ percentile), for the balloon and the fingertip.

**Figure 5.**
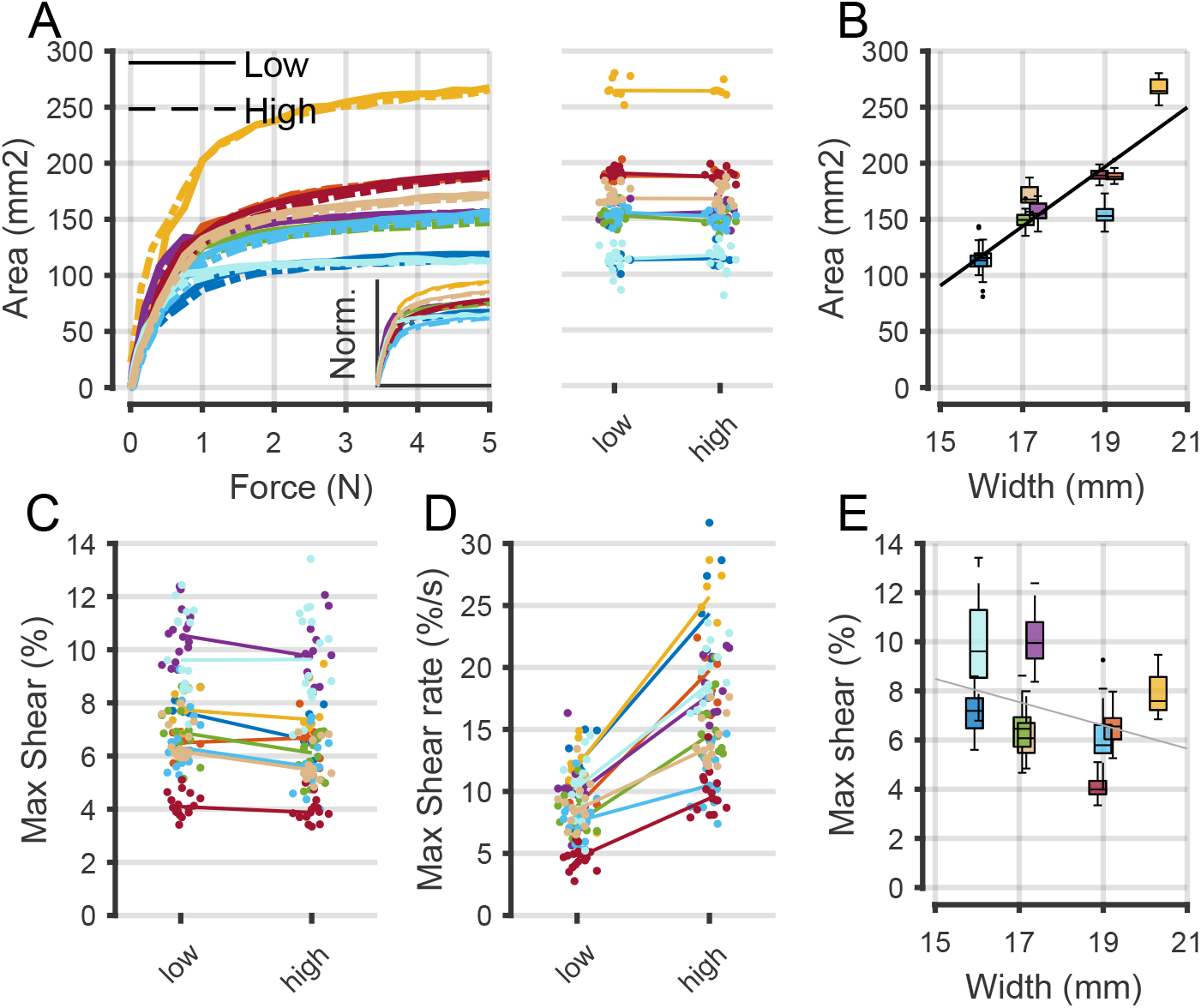
Individual variability, and the effect of loading speed and fingertip size. (**A**) Contact area in the two different speed conditions. Left: evolution versus normal force for each subject (different colors correspond to each subject), averaged across trials, and normalized to fingertip width squared in small panel. Right: final contact area at 5N in all trials. (**B**) Contact area at 5N versus fingertip width. (**C**) Max shear strains at 5N in each speed condition in all trials. (**D**) Max shear rate during loading (at 0.5N) in each speed condition in all trials. (**E**) Max shear at 5N versus fingertip width. Maximum values of shear total and rate were computed as 95^*th*^ percentile. Black fit given in black when fit is significant (<0.05) and light gray when non-significant.

Despite these similarities to the balloon, major differences in mechanical response were evident. For instance, the fingertip contact area grows rapidly and saturates at around 1-2N whereas the balloon’s contact area increases linearly with normal force (Fig. 4F). Additionally, the spatial extent of the deformation front was analyzed relative to the meridional angle (Fig. 4G). The fingertip’s deformation front remains localized near and slightly outside the contact border, whereas the balloon’s deformation spreads significantly into the periphery as soon as contact begins. Finally, the circumferential strain in the balloon evolves linearly and does not saturate, unlike the fingertip (Fig. 4H). Notably, the fingertip exhibits an inflection point around 1-2N (consistent with previous findings by B. Delhaye et al. (2014)), yet deformation continues even after the contact area saturates. These differences highlight the unique biomechanical properties of the fingertip, including its nonlinear response to loading and its capacity for localized deformation.

### Effect of loading speed, fingertip size, and differences across participants

Next, we studied the effect of the loading speed on the fingertip response on the contact area and the deformation. Although the contact area largely differed in amplitude across subjects (Fig. 5A), all traces showed a consistent inflection point around 1N and followed a similar growth pattern during normal loading when normalized to fingertip size (Fig. 5A bottom). Statistical analysis showed that the contact area was not significantly affected by loading speed at 5N (t_229_ = −1.83; p=0.07). To investigate the variability in contact area across participants, we analyzed the effect of fingertip size. As expected, contact area variability was greatly impacted by fingertip width (t_229_ = 5.5247; p=8.94e-8). We also observed substantial variability of total deformation and deformation rate across subjects at both speeds (Fig. 5B-C). Notably, inter-trial variability was present in all subjects, with average shear rate standard deviations of 1.92 %/s and 2.69 %/s for low- and high-speed conditions, respectively. Statistical analysis revealed a significant effect of speed on total shear at 5N (t_229_ = −4.37; p= 1.9e-5), although the observed ratio of 0.92 (mean) between speeds suggests minimal viscous effects. Finally, we found an important increase in the shear rate during loading, with a ratio of 1.94 *±* 0.53 (median *±* 95% CI) (Fig. 5C). This ratio closely aligns with the observed loading rate ratio (1.93 *±* 0.125; median *±* 95% CI). To further explore inter-subject differences in strain, we examined the effect of fingertip size (measured once per subject) on shear strain. While a trend was observed, the effect was not statistically significant (t_229_ = −1.2041; p=0.23).

### Friction role during contact initiation

Finally, we examined whether friction affects the deformation observed during fingertip loading (Fig. 6). Indeed, recent work has shown a significant effect of friction on the displacement field of the finger skin at the contact interface, linking the ability to discriminate large frictional differences to these differences in displacement (Willemet et al. 2021). Here, despite the small range of variation in friction (*∼*0.5 for each subject), the dynamic coefficient of friction significantly affected skin displacement in the contact area, specifically partial slippage during contact initiation (see Material and Methods, t_227_ = −3.173; p = 0.0017). A linear mixed-effects model provided a strong fit to the data, with an adjusted R^2^ = 0.7440. Loading speed also significantly influenced partial slips (t_227_ = 4.0389; p < 10-4), while the direction of sliding did not have a significant effect (t_227_ = −0.0732; p = 0.94). Additionally, as we controlled for the angular displacement of the index finger with non-null attack angle, a non-negligible residual tangential force (TF) was generated at the interface. At a normal force of 5N, this residual TF was 2.17N *±* 0.45N (median *±* 95% CI). The residual TF has a significant effect on the partial slip (t_227_ = −5.2639; p = 0.0226). These findings suggest that indeed, friction strongly influences partial slips during initial loading and as such, could serve as a tactile cue for friction discrimination and grip adjustment (Willemet et al. 2021; Willemet et al. 2024; Schiltz et al. 2022).

**Figure 6.**
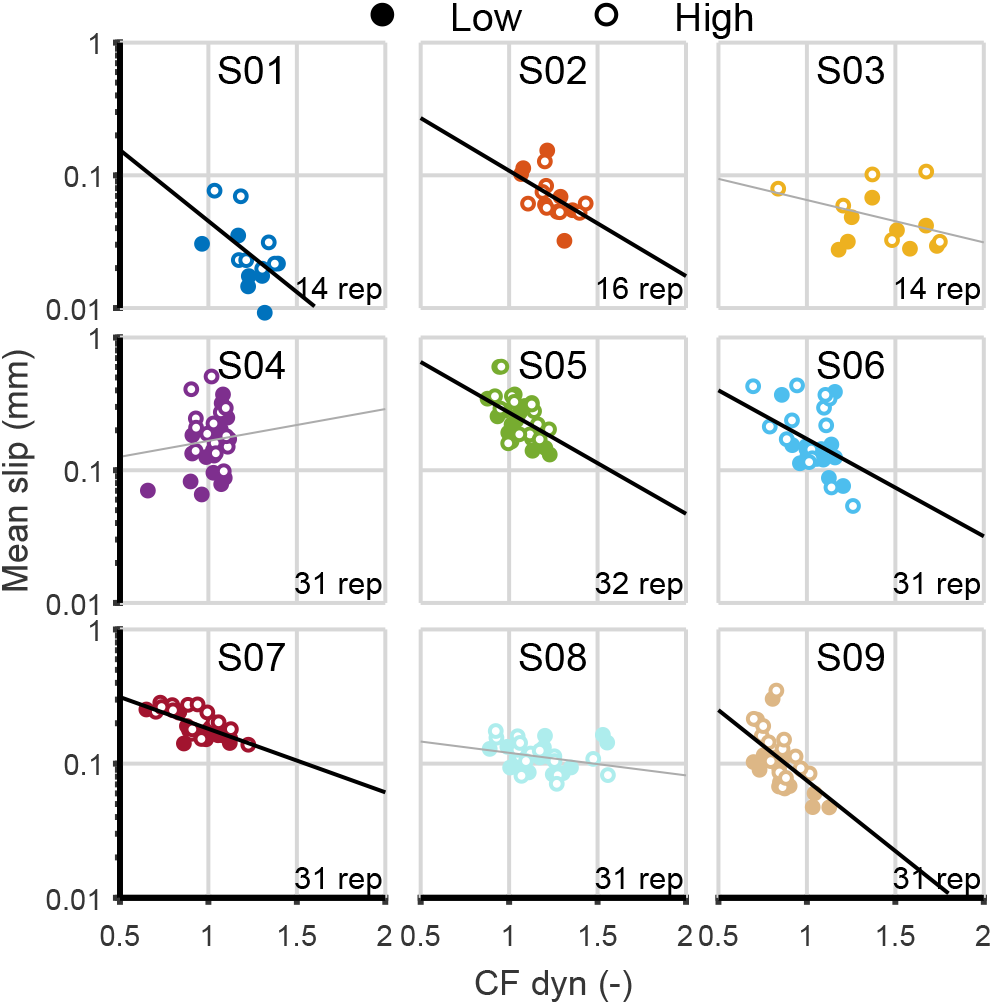
Friction role on the contact initiation. Each plot shows the relation between the mean slip computed during the loading and the coefficient of dynamic friction computed following the loading. Plain dots referred to low-speed condition and open dots to high-speed condition. The linear fit was done over the mean slip in a log scale (uniformly distributed residuals) and is plotted in black when the fit is significant (<0.05) and light gray when not significant.

## Discussion

This study represents, to our knowledge, the first empirical measurement of local strain across the entire glabrous side of human fingertip during natural object interaction. Leveraging an advanced imaging approach to reconstruct the fingertip’s 3D surface, we successfully measured skin deformation patterns when normally loaded onto a glass plate. Our results revealed significant surface strain amplitudes both inside and outside the contact area. We observed strain waves propagating from the initial point of contact toward the periphery in the meridional direction, closely following the contact border as the loading increased. Notably, as soon as contact was initiated (<0.05N), a peak strain rate was detected, highlighting the skin’s remarkable tactile softness at low forces. While strain amplitudes varied significantly across individuals, the wavefront pattern was consistent across all participants.

### 3-D approach benefits

A single-camera setup imaging the fingertip from below requires projecting the 3-D curvature of the fingertip onto a 2-D plane. This projection, if unaccounted for, compromises measurement accuracy and may result in approximations, particularly during loading, when the fingertip undergoes substantial shape deformation (Willemet et al. 2021). In contrast, the 3-D approach used in this study preserves the fidelity of the measurements and provides a more accurate representation of the fingertip’s deformation dynamics.

### Strain amplitude and localization

During loading, the fingertip skin experienced contractive strain in the meridional direction and primarily elongation in the circumferential direction. We compared these principal strains with those predicted by a simplified structural model of the fingertip, specifically a membrane model (Srinivasan 1989). Although this model effectively captured deflection profiles under line load, it grossly approximated surface strains and failed to accurately reflect the evolution of contact area and strain amplitudes with force. These discrepancies highlight the need for additional biomechanical properties, such as non-null bulk shear modulus and anisotropy of collagen fibers, to better characterize the fingertip’s response during even simple interactions.

Our results showed that the amplitude of fingertip skin strain scaled with the applied normal force, ranging from 5% to 10% at a load of 5N. Besides, these values are much lower than the amplitudes observed during slip, which reach up to 50% (B. Delhaye et al. 2016; B. P. Delhaye et al. 2021).

Furthermore, we show that strain rates peaked at very low forces (< 0.05N) and progressively decreased as the loading force increased. This decrease at low forces aligns with the interpretation of Serina et al. (1998), who attributed it to the convex curvature of the fingertip, suggesting a profound effect of fingertip geometry on its deformation behavior under load (Srinivasan and Dandekar 1996).

In addition, the localization of the strain wavefront provides valuable insight into the extent of the contact area, as it consistently progressed along with and slightly ahead of the contact boundary. This resulted in a total meridional strain that accurately delimited the contact area.

### Dominant Elastic behavior

In quasi-static compression conditions comparable to those in this study, previous research (Pawluk and Howe 1999; Jindrich et al. 2003) showed that fingertip force response is minimally affected by loading speed. Wiertlewski and Hayward (2012) further demonstrated that the bulk mechanical properties of the fingertip can be reliably characterized as elastic up to frequencies of 100 Hz. Consistent with these findings, our measurements of the whole skin surface response confirm that under loading speeds relevant to object manipulation, fingertip surface strain predominantly exhibits elastic mechanical properties, with little evidence of viscous effects.

### Variability

Our findings revealed notable individual variability in both contact area and strain amplitude (Dunilac et al. 2023). While the variability in contact area is well explained by differences in fingertip size, skin strains showed only a weak trend and were not significantly affected. This lack of correlation is likely due to inter-individual differences in skin elasticity, which can be influenced by factors such as stratum corneum thickness, skin moisture content, and daily tactile experiences (Wang and Hayward 2007).

We also observed inter-trial variability above the measurement noise level, suggesting biomechanical changes in skin between trials. These changes could be due to occlusion and sweat rates, as moisture levels are known to fluctuate significantly during object manipulation (André et al. 2010). Variability in the frictional properties at the contact interface further underscores these biomechanical differences.

### Friction effect

Leveraging the observed inter-trial variability in friction, we also show that natural initial contact could serve as a cue for discriminating friction. Indeed, partial slip at the contact interface correlated with the dynamic coefficient of friction measured during subsequent sliding events. This result confirms recent results by Willemet et al. (2021), which used a 2D imaging setup coupled with an ultrasonic platform to show similar relationships. We identified a significant effect of loading speed on partial slip, which may be attributed to stiffer skin dynamics caused by viscoelasticity. However, it is also possible that the observed speed effect reflects an increase in measurement noise, as higher loading speeds were associated with greater noise levels.

### Limitations

As a “carrier of deformation information” (Dong and Pan 2017), the speckle pattern is a key aspect of the DIC approach and needs to be carefully fabricated to ensure accurate measurements. In this study, we observed localized fading of the speckle pattern, particularly on the fingertip ridges, due to friction. While this fading could potentially impact the method ‘s precision, no significant increase in noise level was observed across trial repetitions, suggesting that the overall data integrity was maintained. Furthermore, the application of ink to create the speckle pattern influenced the frictional properties of the fingertip, as evidenced by variations in measured friction. However, it is unlikely that the biomechanical properties of the skin were significantly affected by it (Kao et al. 2022). Finally, analyses of surface strains were only made on the distal end of the fingertip. Expanding the field of view to include more of the fingertip would enhance measurement coverage and improve precision by increasing the overlap between stereo pairs of cameras. Such improvements could provide a more comprehensive representation of fingertip deformation during interactions.

### Perspectives

The field of soft robotics has garnered significant attention in recent years, particularly in biomimetic soft robotics (Dong et al. 2022). Efforts have been made to develop efficient soft robotic fingertips, often employing membrane models to mimic human fingertip mechanics (McInroe et al. 2018; Shi et al. 2022). Here, we show gross similarities in local strain patterns between human and such robotic fingertips. Such empirical measurements could provide valuable insights for designing biomimetic soft tactile sensors, such as those incorporating electronic skin (Yang et al. 2019), which aim to replicate human touch perception by effectively extracting tactile cues.

Interestingly, evidence has shown that fingertip size (Peters et al. 2009) and skin stiffness (Li and Gerling 2023) influence tactile acuity. Hence, future studies could combine psychophysical experiments with strain measurements to investigate the contribution of skin elasticity – reflected by the amplitude of skin local deformation – to human tactile acuity. The application of the DIC approach is also relevant in the case of ultrasonic friction-modulation surfaces. This technique modulates friction by creating small cushions of air between the fingertip and the vibrating surface. Optical imaging methods like FTIR often experience reduced image contrast at high vibration frequencies, compromising measurement quality (Wiertlewski et al. 2016; Bochereau et al. 2017). Alternatively, the DIC method, which uses an artificial speckle pattern, is immune to such issues, offering a more robust solution for reconstructing deformation under such conditions.

From a neurophysiological standpoint, recordings show that the first spike latency and response strength of tactile afferents of type 1 (SA-I and FA-I) are related to their receptive field location on the fingertip (Birznieks et al. 2001; Johansson and Birznieks 2004). Strains measured here — acting as the mechanical driver for afferent responses — are consistent with these findings, with a wavefront (strain rate) originating at the center of contact and progressing to the periphery while decreasing in amplitude. Evidence also indicates that afferents are particularly responsive at the border of contact (Bisley et al. 2000), which corresponds to the region where the highest concentration of strain (total strain) across the fingertip surface is observed. Therefore, integrating the proposed measurement strategy with microneurography recordings would provide a powerful framework to confirm those observations and accurately model individual tactile afferent responses to skin deformation (B. P. Delhaye et al. 2021). This combined approach could, in turn, help extend the capabilities of existing frameworks like TouchSim to simulate a broader and more realistic spectrum of tactile stimuli (Saal et al. 2017).

## Conclusions

Everyday tactile interactions involve complex force dynamics, including tangential loading of the fingertip. The extent to which remote tactile receptors can convey information about localized and transient finger-object slips remains an open question. To address this, we intend to use the methods described in this study to investigate the fingertip’s biomechanical response during slippage under shear stimulation. In conclusion, by translating tactile interactions into local deformation patterns, this tool will help to characterize the first stage of processing of tactile stimuli as it ascends the somatosensory pathway.

## Supplementary Materials

Supplemental Movie M1 to M4 (link).

## Acknowledgments

We are grateful for the technical assistance of Arnaud Browet in the MultiDIC toolbox adaptation.

## Funding Statement

This work was supported by a grant from the European Space Agency, Prodex (BELSPO, Belgian Federal Government). BD is mandated by the Belgian National Fund for Scientific Research (FSR-FNRS).

## Notes

https://osf.io/afdmh/?view_only=87b679e4bca64b19b7465e329b793cde

### Competing Interest Statement

The authors have declared no competing interest.

### Summary of Updates

Revisions were made to the Introduction and Discussion sections from a neurophysiological perspective to better clarify the implications of the results. A paragraph was added in memory of Vincent Hayward's work. Minor typographical errors were also corrected.

